# Quantitative *in vivo* evaluation of the reverse β-oxidation pathway for fatty acid production in *Saccharomyces cerevisiae*

**DOI:** 10.1101/201616

**Authors:** Paulo Gonçalves Teixeira, Verena Siewers, Jens Nielsen

## Abstract

Production of fatty acids using engineered *Saccharomyces cerevisiae* cells is a challenging task in part due to low efficiency of the native fatty acid biosynthesis pathway. One option for improving production efficiency relies on exploring alternative fatty acid production pathways with either improved kinetics, thermodynamics or yield properties.

In this work, we explored the reverse β-oxidation pathway as an alternative pathway for free fatty acid production. Different gene combinations and analysis methods were tested for assessing pathway efficiency when expressed in the yeast *Saccharomyces cerevisiae*. Even though different alternatives were tested, quantitative analysis showed no improvement or major change in fatty acid production of the tested strains in our conditions. This lack of improvement suggests that the tested pathway designs and constructs are either nonfunctional in the tested conditions or the resulting strains lack a metabolic driving force that is needed for a functional pathway.

We conclude that expression of the reverse β-oxidation pathway in *S. cerevisiae* poses many challenges when compared to expression in bacterial systems. These factors gravely hinder development efforts and success rate for producing fatty acids through this pathway.

## Introduction

Fatty acids are an important precursor for synthesis of sustainable replacements for liquid transportation fuels and oleochemicals (Lennen and Pfleger, 2013). Oleochemicals are chemicals commonly obtained and derived from plant oils. However, production of oils from plant biomass has several drawbacks such as competition with fertile land for food and difficulty in optimization and tailoring of desired oil compositions. Due to this, production of fatty acids using engineered microbial cells provides a promising alternative with numerous applications and advantages for sustainable development (Nielsen and Keasling, 2016).

Although there are several microorganisms with high native fatty acid production capabilities, the use of model microorganisms such as the yeast *Saccharomyces cerevisiae* benefits from a variety of tools for straightforward genetic and metabolic engineering and a vast knowledge on its metabolism and cellular functions. This opens doors towards creating and improving strains for production of tailored chemicals (Ostergaard et al., 2000).

Fatty acid production in yeast is mainly performed by a type I fatty acid synthase (FAS) system, characterized by the organization of their catalytic domains where the eight FAS component functions are distributed among only 2 polypeptides encoded by the genes FAS1 and FAS2. The FAS complex catalyzes all the activities necessary for fatty acid synthesis from acetyl-CoA and malonyl-CoA (Schweizer and Hofmann, 2004; Tehlivets et al., 2007). In the FAS complex, acyl chains are bound to an ACP domain and go through all the reduction steps catalyzed by the different protein domains. The FAS complex uses malonyl-CoA as elongation building block, elongating the fatty acid acyl chain by two carbons for each malonyl-CoA consumed. When chain elongation is finished, the acyl-ACP is converted to acyl-CoA and released in this form to the cytosol (Lomakin et al., 2007; Zhu et al., 2017).

An alternative FAS system, mainly present in bacteria and archaea, is the type II FAS. In this system, the fatty acid biosynthesis machinery is composed of discrete monofunctional enzymes. (Schweizer and Hofmann, 2004; White et al., 2005). In eukaryotic mitochondria, also FAS systems resembling the type II systems are found, which are usually responsible for synthesis of specialized fatty acids required for certain cell functions, like lipoylation. These systems are separate and independent from the cell’s main FAS complex (Hiltunen et al., 2009).

Even though type I, type II and mitochondrial FAS systems use ACP as an acyl-binding domain for fatty acid biosynthesis, a mitochondrial pathway was identified in *Euglena gracilis* that uses coenzyme A (CoA) as a binding cofactor for fatty acid synthesis (Inui et al., 1984) and acetyl-CoA instead of malonyl-CoA as elongation unit. The reactions involved in this fatty acid biosynthesis pathway are very similar to the reactions in the β-oxidation fatty acid degradation pathway if these occurred in the reverse direction. Therefore, this pathway was described already in 1984 as the “reverse β-oxidation” (Inui et al., 1984). The main difference to the activities existent in β-oxidation enzymes is the presence of a trans-2-enoyl-CoA reductase, which was characterized as a novel family of enzymes that used NADH instead of NADPH as part of a proposed fatty acid biosynthesis pathway (Hoffmeister et al., 2005). Presence of this enzyme was also posteriorly detected in *Treponema denticola* and its activity characterized (Tucci and Martin, 2007).

Production of fatty acids in *Saccharomyces cerevisiae* through the FAS complex is a tightly regulated process involving regulation of both precursor and active enzyme levels (Chirala, 1992; Hasslacher et al., 1993; Schüller et al., 1994). For this reason, improving its efficiency for applications in production of fatty acids might be a challenging task, bringing the need for exploring alternative pathways that allow for efficient fatty acid production in *S. cerevisiae*.

The reverse β-oxidation pathway provides an alternative for fatty acid production. By using acetyl-CoA instead of malonyl-CoA as an elongation building block, the reverse β-oxidation is a more energetically efficient pathway for fatty acid production, since the conversion of acetyl-CoA to malonyl-CoA requires use of one ATP molecule. Furthermore, this pathway uses NADH as reducing factor instead of NADPH, which is an advantage in *S. cerevisiae* which has evolved for high cytosolic NADH turnover during fermentation of glucose (Bakker et al., 2001).

The reverse β-oxidation pathway was successfully employed, further optimized and expanded in *E. coli* for production of short and long chain n-alcohols, fatty acids and 3-hydroxy-, 3-keto- and trans-D2-carboxylic acids using either endogenous or heterologous genes (Clomburg et al., 2012; Dellomonaco et al., 2011).

A proof of concept on expressing a reverse β-oxidation pathway in *S. cerevisiae* for formation of butanol and short chain fatty acids has also been previously reported (Lian and Zhao, 2015), but findings reported here pose doubts on the validity of this study. However, quantitative data for calculation of pathway efficiencies for free fatty acid production have not been properly provided. Furthermore, the effect of different strategies and gene combinations has also not been extensively explored in previous works.

In this study, we first explore some theoretical and conceptual properties of this pathway. Then, we aimed to express different gene combinations in *S. cerevisiae* in order to assess the feasibility of the pathway with supporting quantitative data. Quantification of fatty acids in the cell is also done both at the level of total cellular fatty acids and with a focus on short chain free fatty acids.

## Results

### Pathway design

A previous reverse β-oxidation design in *E. coli* (Clomburg et al., 2012) was used as a reference for rational construction of the pathway in yeast. Since this pathway uses acetyl-CoA as building block and NADH as reducing factor, we proposed to express the pathway in the cytoplasm due to substrate and cofactor availability.

To better understand the functionality of the pathway, we conceptualized the reverse β-oxidation pathway as composed of 3 main modules as shown in **Figure 1**: Initiation, elongation and termination.

**Figure 1.**
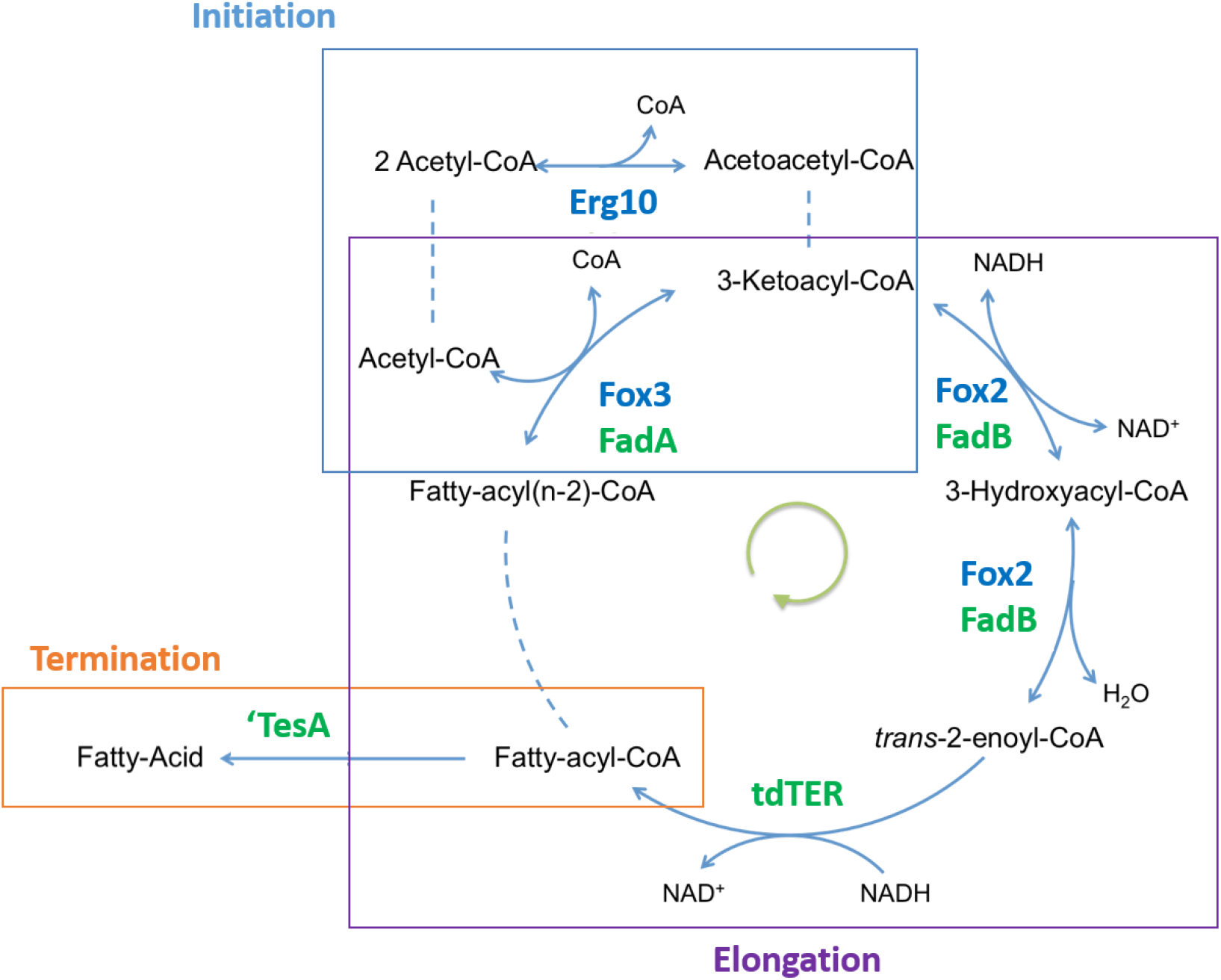
Schematic representation of the reactions involved on the reverse β-oxidation pathway. The squares indicates the initiation strategy for formation of acetoacetyl-CoA using Erg10 (acetyl-CoA acetyltransferase) for the condensation of 2 acetyl-CoAs. Following that, the pathway is composed of Fox3/FadA (3-ketoacyl-CoA thiolase), Fox2/fadB 3-hydroxyacyl-CoA dehydrogenase/enoyl-CoA hydratase multifunctional enzyme) and tdTER (trans-2-enoyl reductase) for elongation. Termination is performed by cleavage of the fatty acyl-CoA into fatty acids by the thioesterase ‘TesA.

#### Initiation

The initiation step is characterized by the formation of acetoacetyl-CoA via condensation of 2 acetyl-CoA. This reaction carries the same biochemistry as the condensation of a longer acyl-CoA with acetyl-CoA, which is a reaction that is naturally present in the reverse direction in the β-oxidation pathway, catalyzed by the 3-ketoacyl-CoA thiolase Fox3 in *S. cerevisiae.* We hypothesized that Fox3 would have a higher affinity for longer acyl-CoAs than for acetyl-CoA, and as such considered alternative enzymes with the adequate substrate specificity. The acetyl-CoA C-acetyltransferase Erg10 from *S. cerevisiae* catalyzes the first step of the mevalonate pathway where 2 acetyl-CoA are converted to acetoacetyl-CoA and as such was selected as the ideal candidate for the initiation step.

Since the thermodynamics of reaction catalyzed by Erg10 has a largely positive ΔG°’ of 26.7 kJ.mol^-1^ and therefore seems thermodynamically unfavourable, we decided to include *NphT7* from *Streptomyces sp. (Okamura et al., 2010)* in order to boost the reaction and increase acetoacetyl-CoA supply for the first cycle to run. However, this was earlier shown in other studies by our lab (Tippmann et al., 2017) to not be an efficient alternative to Erg10 when used *in vivo* to catalyze this reaction and was eventually abandoned.

#### Elongation

Elongation is the process by which the acyl chain in acyl-CoA is increased by 2 carbons using acetyl-CoA as a building block. This cycle is composed of 4 reactions, in which the first 3 reactions are present in β-oxidation in the reverse direction. Since the β-oxidation enzymes involved in this process are reversible, we used either the native *S. cerevisiae* β-oxidation enzymes 3-ketoacyl-CoA thiolase Fox3 and the 3-hydroxyacyl-CoA dehydrogenase/enoyl-CoA hydratase multifunctional enzyme Fox2 or their *E. coli* homologues FadA and FadB. Since β-oxidation in yeast is a peroxisomal process, the peroxisome targeting signals were identified using SignalP and removed in order to express a cytoplasmic form of the proteins (cFox3 and cFox2 respectively).

The first reaction in β-oxidation, catalyzed by Pox1, is an irreversible and very thermodynamically favorable reaction with a ΔG°’ of −42.6 (Table 1) that uses O_2_ as electron acceptor for oxidation of the acyl-CoA molecule. To have a functional reversion of the β-oxidation and allow a thermodynamically feasible production of fatty acyl-CoA, *POX1* was deleted and a trans-enoyl reductase was expressed to catalyze a favorable reaction in the reverse direction. For this, tdTER from *Treponema denticola* (*Tucci and Martin, 2007*) was chosen as candidate due to its efficiency, substrate specificity and reported irreversibility of the reaction in the reductive direction. By expressing *tdTER*, we allow the reaction to occur in the opposite direction, using an irreversible enzyme that uses NADH as the cofactor in the reduction of the trans-enoyl-CoA to acyl-CoA. As shown in **Table 1**, this reaction influences drastically the thermodynamics of the pathway.

**Table 1.** Calculated ΔG°’ values for the reverse β-oxidation pathway.

| Enzyme | Reaction<sup>1</sup> | $\Delta G^{\circ'}$<br>(kJ mol<sup>-1</sup>) |
| --- | --- | --- |
| Erg10/Fox3/YqeF/FadA | 2 acetyl-CoA → acetoacetyl-CoA + CoA | 26.7 |
| Fox2 | 3-oxoacyl-CoA + NADH + H<sup>+</sup> → hydroxyacyl-CoA + NAD<sup>+</sup> | -19.9 |
| Fox2 | hydroxyacyl-CoA → *trans*-2-enoyl-CoA + H<sub>2</sub>O | 2.1 |
| tdTER | *trans*-2-enoyl-CoA + NADH + H<sup>+</sup> → acyl-CoA + NAD<sup>+</sup> | -57.9 |
| TesA | acyl-CoA + H<sub>2</sub>O → fatty acid + CoA + H<sup>+</sup> | -35.5 |
| Pox1<br>(oxidative reaction)<sup>2</sup> | acyl-CoA + O<sub>2</sub> → *trans*-2-enoyl-CoA + H<sub>2</sub>O<sub>2</sub> | -42.6 |

<sup>1</sup> Reactions are presented and $\Delta G^{\circ'}$ s were calculated in the synthetic (reductive) direction (except for POX1 reaction which is presented in the oxidative direction).
<sup>2</sup> The reaction for Pox1 is shown separately in the last column since Pox1 is not part of the reverse β-oxidation pathway and shown here to illustrate how it influences the thermodynamics of the β-oxidation pathway in native conditions.

#### Termination

In order to efficiently produce fatty acids in *S. cerevisiae*, acyl-CoAs need to be pulled from the elongation cycle and converted either into free fatty acids or fatty acid-derived chemicals. As a proof of principle, we expressed the *E. coli* truncated thioesterase ‘TesA (Cho and Cronan, 1993), a broad specificity thioesterase that is able to cleave acyl-CoAs of different chain lengths to their correspondent free fatty acids. This would allow us to check for production capacity of the pathway for different chain length products, especially shorter chain fatty acids.

### Thermodynamics and NADH considerations

In order to evaluate the feasibility of the pathway, it is important to check parameters such as thermodynamic barriers and balance of precursors and cofactors. Using the additive group contribution method, the ΔG°’ values of the reactions involved were calculated. **Table 1** shows the values obtained.

A first look at the ΔG” values shows that the reaction catalyzed by Pox1 is one of the determinant steps in the β-oxidation pathway of fatty acids in terms of thermodynamic feasibility due to a highly negative ΔG” in the oxidative direction. It is also evident that tdTER has a very favorable ΔG” value in the synthetic (reducing) direction, which explains how replacing the reaction catalyzed by Pox1 with the reaction from tdTER can change the overall thermodynamic feasibility of this pathway for fatty acid synthesis.

ΔG” values account for the Gibbs free energy at equal concentrations of all the molecules involved, not considering that in an intracellular environment these components can vary by several orders of magnitude. As it is, the pathway seems thermodynamically feasible. We can consider that the main thermodynamically challenging step would be the first reaction, catalyzed by Erg10/Fox3/FadA. However, in *S. cerevisiae* we observe flux through Erg10 upon formation of acetoacetyl-CoA for the mevalonate pathway. So we can argue for some supply of acetoacetyl-CoA to the reactions catalyzed by Fox2, even though other enzymes compete for the same precursor and thus limit its availability. Concerning the Fox2-catalyzed reaction, we can consider most intermediates virtually non-existent in the cytosol as a starting condition, which makes the reaction thermodynamics dependent on the NADH/NAD+ ratio. Solving the Gibbs free energy equation to find K_eq_ in an equilibrium situation (ΔG=0) and assuming the concentration of both ketoacyl-CoA and hydroxyacyl-CoA is equal (so that K_eq_=[NAD+]/[NADH], then the reaction would be feasible unless the NAD+/NADH ratio would be higher than 2.7×10^3^, which we can safely assumed not to be the case in standard intracellular growth conditions in yeast (Bekers et al., 2015).

In terms of cofactor balance, the pathway is balanced for NADH, since for each cytosolic acetyl-CoA formed from ½ Glucose, 2 NADH are generated (considering the main native acetyl-CoA generating pathway in yeast to be the PDH bypass through pyruvate → acetaldehyde → acetate → acetyl-CoA), which is also how much is used by the pathway per elongation cycle in which 1 acetyl-CoA is incorporated (pathway stoichiometry shown in **Table 2**). The 2 extra NADH generated together with the first acetyl-CoA could easily be used by reducing the final fatty acid molecule to a fatty alcohol or put towards ATP generation. The major stoichiometric problem is that to form acetyl-CoA through this route, the ATP balance is negative, which leads to the requirement of some NADH oxidation through mitochondrial respiration to generate enough ATP, this way harming maximum theoretical yields. However, the pathway is much more favorable than fatty acid production through the native FAS system in terms of ATP balance, since the FAS system requires 1 malonyl-CoA per elongation cycle instead of 1 acetyl-CoA, which increases the total cost of the elongation cycle by 1 ATP.

**Table 2.** Stoichiometry of pathways for acetyl-CoA formation and consumption through the reverse β-oxidation compared with the native fatty acid biosynthesis.

| Pathway | Reaction Stoichiometry |
| --- | --- |
| Native yeast PDH bypass pathway <sup>1</sup> | $\frac{1}{2} \text{ glucose} + 2 \text{ NAD}^+ + \text{ATP} + \text{CoA} + \text{H}_2\text{O} \rightarrow \text{acetyl-CoA} + 2 \text{ NADH} + 2\text{H}^+ + \text{CO}_2 + \text{ADP} + \text{Pi}$ |
| Reverse β-oxidation Elongation | $\text{acyl}(n-2)\text{-CoA} + \text{acetyl-CoA} + 2 \text{ NADH} + 2 \text{ H}^+ \rightarrow \text{acyl-CoA} + \text{CoA} + 2 \text{ NAD}^+ + \text{H}_2\text{O}$ |
| FAS Elongation | $\text{acyl}(n-2)\text{-ACP} + \text{acetyl-CoA} + \text{ATP} + 2 \text{ NADPH} + \text{HCO}_3^- \rightarrow \text{acyl-ACP} + \text{CoA} + 2 \text{ NADP}^+ + \text{H}_2\text{O} + \text{ADP} + \text{Pi}$ |

<sup>1</sup> as described in (van Rossum et al., 2016)

### Effect of expressing the reverse β-oxidation pathway on total fatty acid levels

On a first assessment of the pathway, Erg10 from *S. cerevisiae* was used as the initiation enzyme. For the elongation module, the *E. coli* genes *fadA* and *fadB*, already shown to be functional in the reductive pathway in their native host, and the heterologous *tdTER* were used in the pathway construction. As an alternative to *fadA* and *fadB*, cytosolic forms of the *S. cerevisiae FOX3* and *FOX2* genes were also used. The thioesterase ‘TesA was used as the termination step. Combinations of 5 genes were cloned into a multi-copy 2μ plasmid with *URA3* auxotrophy marker, generating 2 different plasmids with alternative constructs of the reverse β-oxidation pathway (pRBye: *ERG10*, *fadA*, *fadB*, *tdTER*, *‘tesA*; and pRByy: *ERG10*, *cFOX3*, *cFOX2*, *tdTER*, *‘tesA*).

For pathway analysis, we used both a wild-type CEN.PK113-11C strain or an engineered strain YJZ06 containing deletions in *POX1*, *FAA1* and *FAA4*. *POX1* catalyzes the first step of the β-oxidation and deletion of this gene is required as stated in the section above. *FAA1* and *FAA4* encode fatty acid synthetases responsible for converting free fatty acids back to acyl-CoA. Deletion of *FAA1* and *FAA4* would avoid potential feedback regulation of formed free fatty acids (FFAs) or reconversion of these into acyl-CoA, which would have a negative effect by reversing the reaction catalyzed by ‘TesA. At this stage, the pRBye plasmid was tested in the wild-type CEN.PK113-11C and both plasmids pRBye and pRByy were tested in YJZ06. For both strains, a plasmid only expressing *‘tesA* (ptesA) was used as a control.

Strains were analyzed for total fatty acids and quantification was focused on the separation of short chain fatty acids since these were the product expected from a functional pathway. However, analyzed strains showed no significant difference in production of short chain fatty acids. No C6 was detected while C8 was at very low levels with only a slight non-significant difference between expressing the pathway or just *‘tesA* as a control (**Figure 2a** and **2b**). In terms of total fatty acid profile and peaks for longer chain fatty acids, no relevant differences were also noticed (example chromatogram shown in **Figure 2c**).

**Figure 2.**
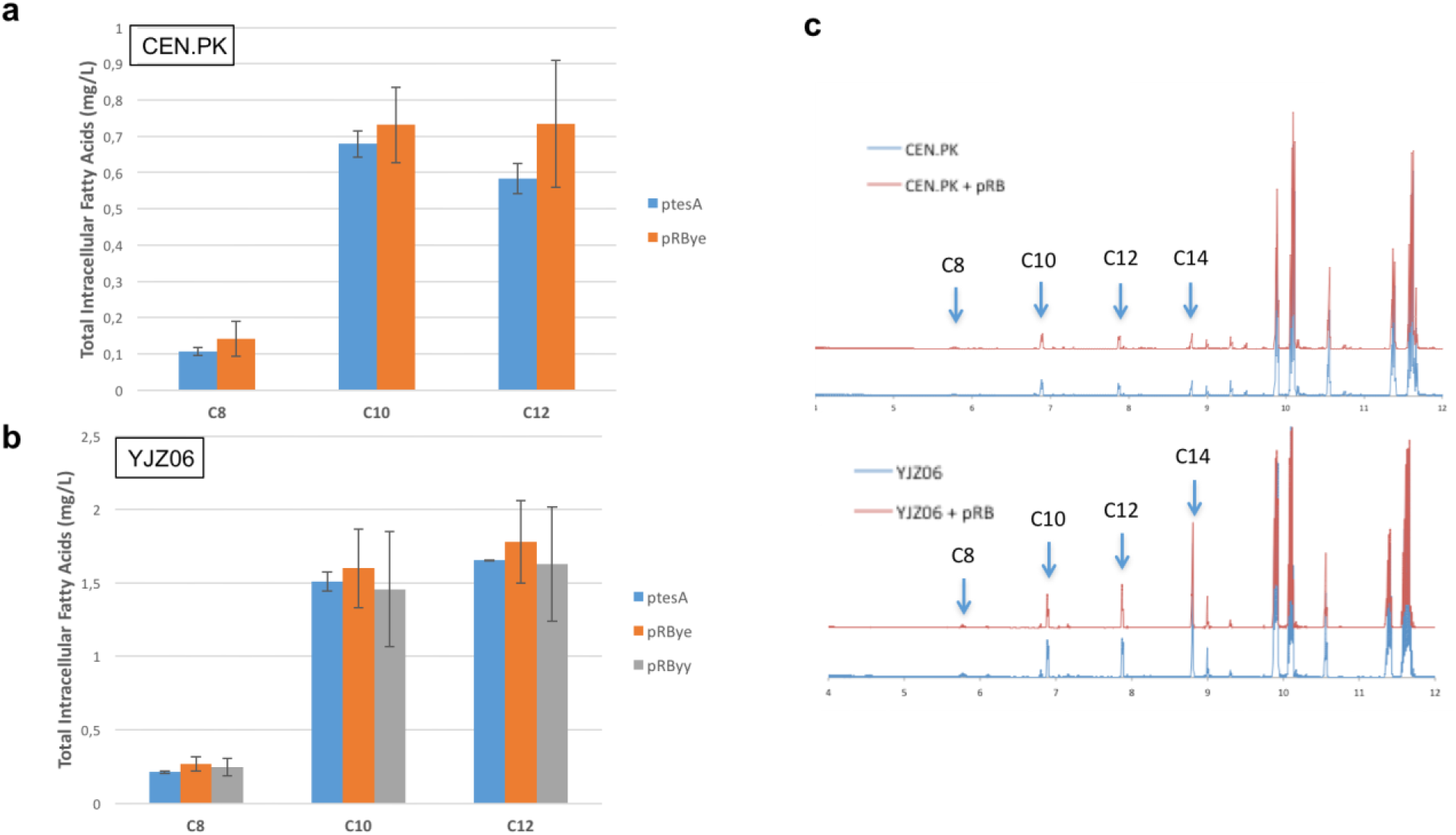
Total intracellular fatty acid levels in strains expressing the reverse β-oxidation pathway. Levels of short chain fatty acids are shown for strains a) CEN.PK 113-11C or b) YJZ06 (CEN.PK 113-11C *Δpox1 Δfaal Δfaa4*) containing a multi-copy plasmid expressing the full reverse β-oxidation pathway (pRBye and pRByy) or only ‘TesA (ptesA). c) Example chromatograms representative of the GC-MS analysis for total fatty acids. Cells were cultured for 72 h in minimal media with 2% glucose. Experiments were performed as biological triplicates and error bars represent standard deviation.

### Alternatives for elongation enzymes

A study on the proof of concept for this pathway in yeast (Lian and Zhao, 2015) showed the inefficiency of both *FOX2*, *fadB and fadA* for catalyzing the reactions required for the reverse β-oxidation when tested from yeast extract using *in vitro* assays. This is in agreement with our previous results. The same study reported successful production of fatty acids through the reverse β-oxidation when using two genes from *Yarrowia lipolytica*, a β-ketoacyl-CoA reductase *YlKR* and a β-hydroxyacyl-CoA dehydratase *YlHTD*, involved in mitochondrial fatty acid synthesis, truncated at the N-terminal to remove the mitochondrial targeting signal. These two enzymes replace the β-oxidation multifunctional enzyme Fox2, catalyzing the two same reactions. Thus, we reconstructed the pathway in a new version expressing *ERG10* for initiation, *cFOX3*, *YlKR*, *YlHTD and tdTER* for elongation, and *‘tesA* for termination. While designing this new version, we identified the existence of a crotonase-like, putative hydroxyacyl-CoA dehydratase, gene in the *Y. lipolytica* genome through sequence comparison with the *YlHTD* gene sequence using BLAST. This gene, here named as *YlCRT* (*Yarrowia lipolytica* crotonase), was included in the design as an alternative to *YlHTD*.

The pathway was tested again by expression from a single 2μ plasmid and was analyzed in YJZ02 (CEN.PK113-11C *Δpox1*) and YJZ06 (CEN.PK113-11C *Δpox1 Δfaa1 Δfaa4*) strains. The strains with the new pathway version were analyzed with a more efficient and sensitive method focused on the detection of extracellular short-chain fatty acids (Zhu et al., 2017).

From the results shown in **Figure 3**, strains expressing the reverse β-oxidation pathway do not have a significant increase in production of short chain fatty acids. At the level of the detected C6 fatty acids there is a slight increase for pRBhtd compared to the control strain, however, this result is not statistically significant for the tested sample groups (p>0.05 using student’s t-Test). Growth rates and final OD values in these strains were also not different between them (data not shown).

**Figure 3.**
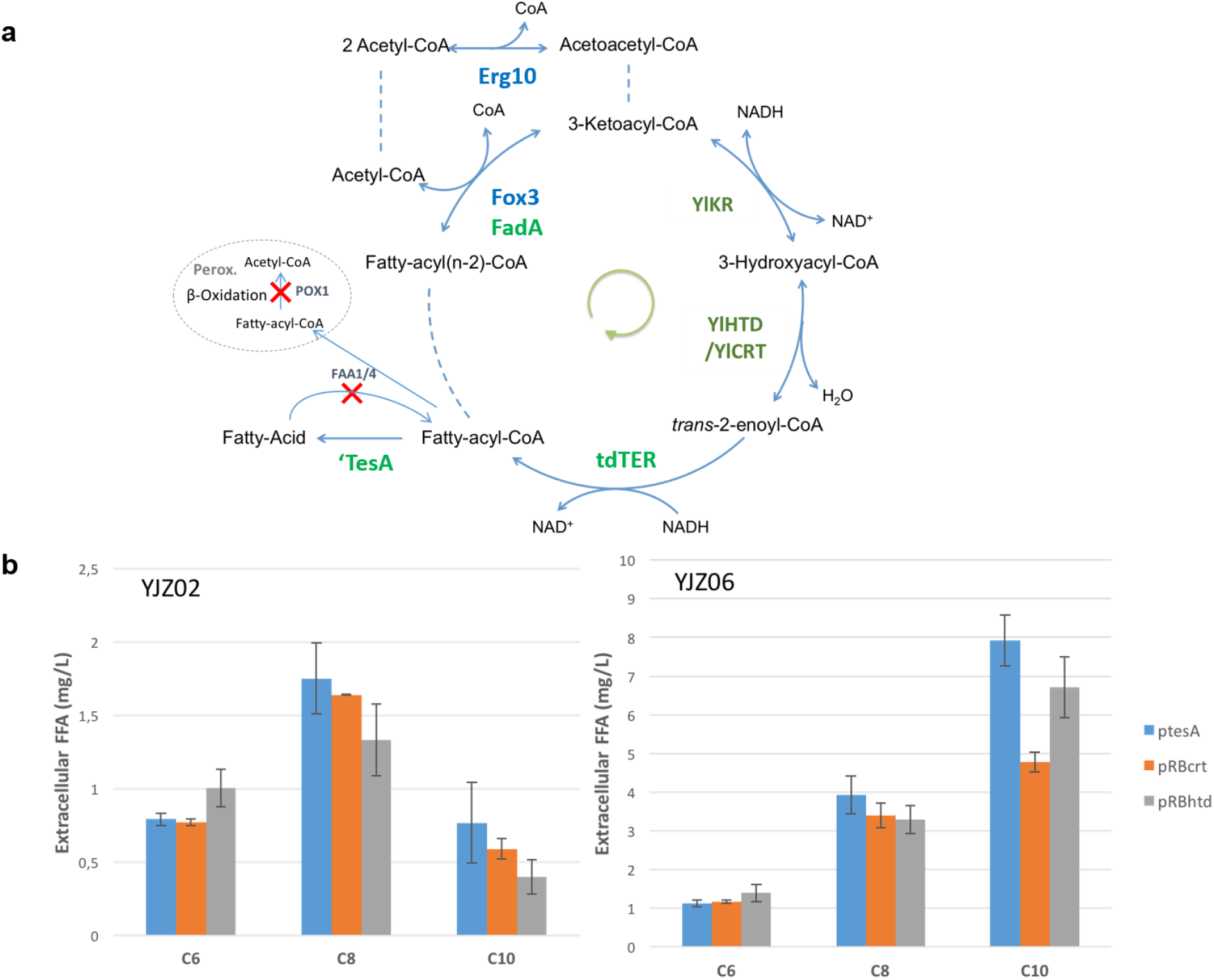
Short chain fatty acids produced from strains expressing the second version of the reverse β-oxidation pathway. a) Schematic representation of the reactions involved on the reverse β-oxidation pathway constructs based on the strategy from Lian & Zhao 2014, expressing YlKR and YlHTD instead of Fox2. An alternative gene to YlHTD, YlCRT was also tested. b) Extracellular short chain fatty acid levels are shown for expression of the pathway in strains YJZ02 (CEN.PK113-11C *Δpox1*) or YJZ06 (CEN.PK113-11C *Δpox1 Δfaal Δfaa4*). The plasmid ptesA containing only *‘tesA* was used as a control, pRBcrt and pRBhtd represents strains with the respective plasmids, expressing the pathway using either YlCRT or YlHTD as alternatives for the hydroxyacyl-CoA dehydratase step. Cells were cultured for 72 h in minimal media with 2% glucose. Experiments were performed as biological triplicates and error bars represent standard deviation.

### Quantitative analysis of previously validated strategies

Even though we replaced Fox2 by the *Y. lipolytica* enzymes, these might not be the only factors influencing the performance of the pathway. Therefore, the original plasmids used in (Lian and Zhao, 2015) were requested from the authors in order to get more insight from own experimentation and quantify the efficiency of the pathway in its functional state. The study uses the cytosolic form of Fox3 (cFOX3) for initiation, this same *cFOX3*, *YlKR*, *YlHTD* and cytosolic Etr1 (*cETR1*) for elongation and the CpFatB1 thioesterase for termination. *cETR1* encodes the endogenous mitochondrial Etr1 without the mitochondrial signal peptide and is used for the enoyl-reductase activity. This gene replaces the function of *tdTER* on the elongation cycle that we used in our previous strategies. CpFatB1 is a thioesterase from *Cuphea palustris* specific for short chain fatty acyl-CoAs, which increases the pathway efficiency in cleaving short chain acyl-CoAs. The system consists of two 2μ plasmids, one containing the reverse β-oxidation genes (rP32 or rP35) with a *URA3* marker and another plasmid containing *CpFatB1* with a *LEU3* marker. These plasmids were introduced in CEN.PK2 and cultured under the same conditions as reported in the reference study (Lian and Zhao, 2015).

Surprisingly, we could again not observe an increase in short chain fatty acids (**Figure 4**). Original strains containing the described plasmids were also requested in order to rule out any possible errors in the transformation or mutated plasmids but with the same result (data not shown).

**Figure 4.**
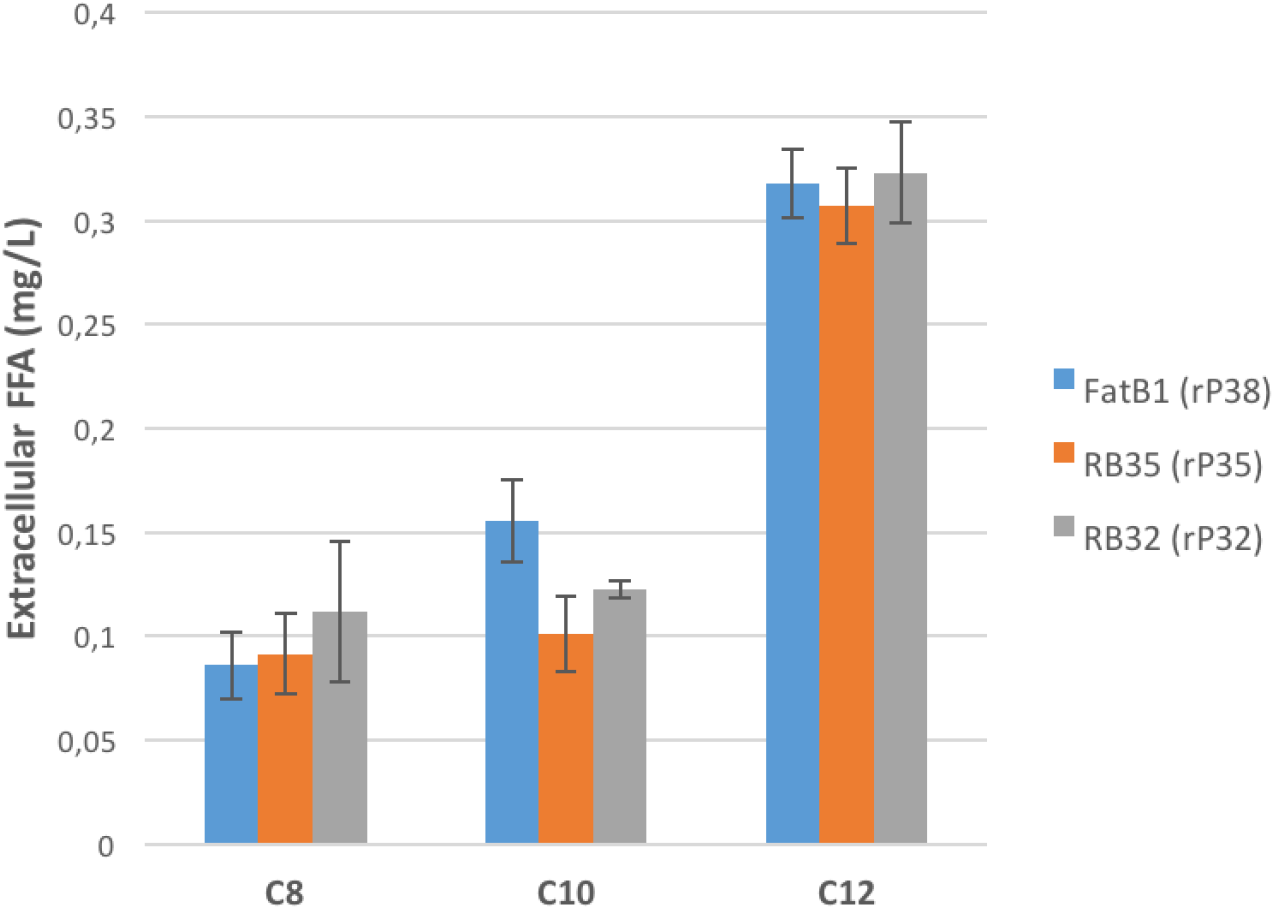
Short chain fatty acids produced from strains transformed with the plasmids from Lian and Zhao, 2014. Short chain fatty acids were measured in supernatants of CEN.PK2 strains transformed with the described rP32, rP35 or rP38 plasmids expressing either a reported functional (rP32 and rP35) or a non-functional (rP38) version of the reverse β-oxidation pathway. All strains were co-transformed with a pCpFATB1 plasmid encoding a thioesterase with affinity towards short chain acyl-CoAs. Cells were cultured as described by Lian and Zhao and experiments were performed as 4 biological replicates, error bars represent the standard deviation.

Many factors can play a role in the pathway activity. Details in media composition, batches, processing and culture conditions might play important roles that will generate unpredictable differences. However, an important note was that, again, a large heterogeneity and instability was observed in these strains containing two different 2μ plasmids. When picked for pre-culture, about 6 colonies for each strain would be selected because half of them would fail to grow on selective media. This effect has been observed before in our lab and we speculate that by using two different 2μ plasmids (therefore sharing the same origin of replication), one of the plasmids could be lost during the process of cell cultivation and create unstable results in different conditions.

## Discussion

There are many challenges in the development of the reverse β-oxidation pathway in yeast. First and foremost, the pathway consists of a set of enzymes ranging from 4 to 6 genes. For many other pathways, one could test the function of each enzyme individually in a metabolic pathway to address its feasibility. In this case however, when dealing with -CoA intermediates, it becomes very difficult to assay and measure these in an easy and straightforward due to low in-vivo concentration and low availability of the specific substrates. (Lian and Zhao, 2015) did measure enzyme activity for different enzymes from cell extracts of all the steps involved, but only for C4 intermediaries and for the first reaction this was assayed only in the reverse direction (acetoacetyl-CoA cleavage). These assays indicate viability for butanol production but there’s still a variable of substrate specificity for chain length that plays an important role for fatty acid production. Therefore, we rely on constructing the complete pathway and accessing the final products for evaluating pathway functionality and efficiency. Being a pathway of 5 to 6 genes, it is time-consuming to investigate performance of individual steps when no changes are detected in the final product.

Considering the hypothesis that a viable solution for pathway assessment exists, we are still confronted with many metabolic engineering challenges. The pathway seems to depend on a high driving force from acetyl-CoA and NADH supply. This is indicated by the need to knock-out fermentative pathways in *E. coli* that would be NADH-consuming (Dellomonaco et al., 2011). Applying this strategy to yeast, (Lian and Zhao, 2015) increased the efficiency of the pathway through increasing NADH supply by simultaneous disruption GPD1 and GPD2 for glycerol biosynthesis and ADH1-ADH4 for ethanol formation. In the same strain, a feedback inhibition-insensitive acetyl-CoA synthetase mutant from *Salmonella enterica* (SeAcs^L641P^) was also overexpressed resulting in improved production of short chain fatty acids through the pathway, supporting the indication that precursor supply is a major limitation in the success of applying the reverse β-oxidation.

Another challenge on this development is how each intermediate is subjected to side reactions. As an example, the acetoacetyl-CoA generated on the first step of the first cycle is used by Erg13 in the mevalonate pathway, which implies already here a direct competition. Furthermore, -CoA intermediates of fatty-acids might be transported through non-specific activities of acyl-CoA transporters to the peroxisome, where Fox2 and Fox3 are still acting in the degradative direction. As far as we know, no studies have been conducted testing activity of these transporters towards these intermediates.

Nevertheless, some of these challenges are also opportunities for development. As examples, restricted performance of a pathway with a need for NADH would allow for adaptive laboratory evolution experiments on a strain without fermentative ethanol metabolism for increasing the efficiency of the pathway as a better NADH sink to improve growth and consequently fatty acid production. A similar study on associating reverse β-oxidation activity with growth in *S. cerevisiae* was previously performed, where the *E. coli* pyruvate dehydrogenase (PDH) complex, a complex responsible for conversion of pyruvate into acetyl-CoA, was expressed in a *Δacs1 Δacs2* yeast strain, which means that cytosolic acetyl-CoA production would be dependent majorly on PDH (Lian and Zhao, 2016). Because PDH needs lipoylation to be active, a lipoylation pathway dependent on octanoic acid production through the the reverse β-oxidation was also expressed. Success of such strategy was highly limited, once again suggesting the weak efficiency of the pathway. However, it lays the foundation for the possibility of development of studies for development of the reverse β-oxidation pathway carrying growth as the screening element.

## Conclusions

Although the reverse β-oxidation pathway is possibly a viable and efficient alternative for fatty acid production in bacteria, it is very challenging to express this pathway in *S. cerevisiae*. We evaluated different possibilities for pathway design and conclude that intrinsic characteristics of the metabolic environment of yeast cytoplasm dramatically hinder the efficiency of this pathway for fatty acid production.

## Author Contributions

PGT, VS and JN conceived the study. PGT carried out the experiments. PGT drafted the manuscript. All authors contributed to the analysis and the discussion of the results. All authors read and approved the final manuscript.

## Acknowledgements

Authors would like to acknowledge Dr. Yongjin Zhou and Dr. Klaas Buijs for helpful discussion and guidance. Parts of this project was funded by the Novo Nordisk Foundation, the Knut and Alice Wallenberg Foundation and the Swedish Foundation for Strategic Research.

## Methods

### Plasmid and strain construction

Endongenous genes *FOX2*, *FOX3*, *ERG10* were amplified by PCR from *S. cerevisiae* CEN.PK113-11C. *Yarrowia lipolytica* genes were amplified from strain A101. Plasmid constructs were generated by fusion PCR and homologous recombination in yeast as reported by (Shao et al., 2009) using pYX212 *(URA3* marker) and p423GPD (*HIS3* marker) as backbones using *S. cerevisiae* CEN.PK113-11C for homologous recombination assembly. Strains and plasmids are listed in **Table 3 and 4**, respectively. A list of primers and sequences for synthetic genes used for plasmid construction are shown in **Table S1 and S2**. Sequence maps for the fragments cloned in pYX212 plasmid for expression of the reverse β-oxidation pathways with corresponding genes, promoters and terminators are schematized in **Figure S2**. *S. cerevisiae* strains were transformed with plasmids and recombination fragments using either lithium acetate and PEG3350 (Gietz and Schiestl, 2007) or electroporation (Thompson et al., 1998).

**Table 3.** *Saccharomyces cerevisiae* strains used in this study.

| Strain | Genotype | Reference |
| --- | --- | --- |
| CEN.PK113-11C | *MATa MAL2-8c SUC2*<br>*his3Δ1 ura3-52* | Euroscarf |
| YJZ02 | *MATa MAL2-8c SUC2*<br>*his3Δ1 ura3-52 pox1Δ* | (Zhou et al., 2016) |
| YJZ06 | *MATa MAL2-8c SUC2*<br>*his3Δ1 ura3-52 pox1Δ*<br>*faa1Δ faa4Δ* | (Zhou et al., 2016) |

**Table 4.** Plasmids used and generated in this study.

| Plasmid | Genotype/Features | Reference |
| --- | --- | --- |
| pYX212 | 2μ multicopy, *AmpR*, *URA3*, *TPI1p*, pYX212t | R&D systems |
| p423GPD | 2μ multicopy, *AmpR*, *HIS3*, *TDH3p*, *CYC1t* | ATCC |
| pRBye | pYX212-*ERG10-fadA-fadB-tdTER-tesA* | This study |
| pRByy | pYX212-*ERG10-cFOX3-cFOX2-tdTER-tesA* | This study |
| ptesA | pYX212-*tesA* | This study |
| pRBhtd | pYX212-*ERG10-cFOX3-cYIKR-cYIHTD-tdTER-tesA* | This study |
| pRBcrt | pYX212-*ERG10-cFOX3-cYIKR-YICRT-tdTER-tesA* | This study |
| rP32 | p426-*cFOX3-cYIKR-cYIHTD-tdTER-EcEutE-CaBdhB* | (Lian and Zhao, 2015) |
| rP35 | p426-*cFOX3-cYIKR-cYIHTD-cETR1-EcEutE-CaBdhB* | (Lian and Zhao, 2015) |
| rP38 | p426-*cFOX3-eGFP-cYIHTD-tdTer-EcEutE-CaBdhB* | (Lian and Zhao, 2015) |
| pCpFatB1 | pH5-*CpFatB1* | (Lian and Zhao, 2015) |

### Growth media

*S. cerevisiae* strains with uracil and histidine auxotrophies were grown on YPD plates containing 20 g/L glucose, 10 g/L yeast extract, 20 g/L peptone from casein and 20 g/L agar. Plasmid carrying strains were grown on selective growth medium containing 6.9 g/L yeast nitrogen base w/o amino acids (Formedium, Hunstanton, UK), 0.77 g/L complete supplement mixture w/o histidine and uracil (Formedium), 20 g/L glucose and 20 g/L agar. Shake flask cultivations were performed in minimal medium containing 20 g/L glucose, 5 g/L (NH_4_)_2_SO_4_, 14.4 g/L KH_2_PO_4_, 0.5 g/L MgSO_4_·7H_2_O. After sterilization, 2 mL/L trace element solution and 1mL/L of vitamin solution were added. The composition of the trace element and vitamin solution has been reported earlier (Verduyn et al., 1992).

### Shake flask cultivations

All experiments were performed with strains cultivated as biological triplicates. This means that three independent transformants were used to start pre-cultures. For these, 3 mL of minimal medium in a 15 mL tube, or 5 mL in a 50 mL tube, were inoculated for the first experiment, and cultivated at 200 rpm and 30°C for 18 h. Subsequently, the pre-culture was used to inoculate 20 mL of minimal medium in a 100 mL shake flask, or 100 mL of minimal medium in a 500 mL shake flask for the first experiment, at an OD600 of 0.1. Shake flasks were incubated at 200 rpm and 30°C for 72 h.

A spectrophotometer (Genesys 20, Thermo Fisher Scientific, Waltham, MA, USA) was used to measure cell growth at designated time points and at the end of the shake flask cultivations. Optical density (OD) was measured by absorbance at 600nm of a diluted culture sample.

### Fatty acid quantification

Initial fatty acid analysis was based on total intracellular fatty acid quantification in the form of fatty acid methyl esters (FAMEs) by extraction and microwave-assisted esterification of yeast biomass according to a previously described method (Khoomrung et al., 2012) with minor modifications. One milliliter hexane, 2 mL boron trifluoride/methanol (14%, Sigma-Aldrich), 10 ug heptanoic acid and 10 ug pentadecanoic acid were added to 10 mg lyophilized biomass for derivatization of total fatty acids. After derivatization, the upper hexane phase was concentrated to about 100 μL, and then analyzed by GC/MS.

Extracellular short chain fatty acids (SCFAs) were extracted and esterified using a previously described method (Leber and Da Silva, 2014) with modifications. Briefly, 0.5 mL 10% (w/v) NaCl, 0.5 mL glacial acetate (containing 10 ug heptanoic acid and 10 ug pentadecanoic acid as internal standards) and 2 mL 1:1 (v/v) chloroform/methanol were added to 4 mL culture broth in extraction tubes (16×100 mm PYREX® culture tubes and GPI 15-415 Threaded Screw Cap, Corning Inc., US). After vortex at 1800 rpm for 30 min, the mixtures were centrifuged at 3000 rpm for 10 min, and the lower chloroform phase was transferred into a clean extraction tube by a glass syringe. FAMEs were generated by mixing 1 mLboron trifluoride/methanol (14%, Sigma-Aldrich) with 200 μL of chloroform extract, and incubation at room temperature overnight. This was based on the ready and fast esterification of free fatty acids by boron trifluoride/methanol The FAMEs were then extracted by adding 1 mL H_2_O and 600 μL hexane, vortexing at 1500 rpm for 10 min, and centrifugation at 1000 g for 10 min. 200 μL taken from the hexane phase were analyzed by GC/MS.

The extracted short/medium chain FAMEs were analyzed by a FOCUS GC/ISQ single quadrupole mass spectrometer system (Thermo Fisher Scientific) using a ZB-50 column (30 m * 0.25 mm * 0.15 μm, Phenomenex). Helium was used as carrier gas (3 mL/min). 1 μL samples were injected (splitless, 240°C), and the oven temperature was set at 30°C for 2 min, increased to 150°C with a ramp rate of 40°C/min, held for 2 min, then increased to 250°C with a ramp rate of 10°C/min, and held for 3 min. The temperatures of MS transfer line and ion source were set as 250°C and 200°C, respectively. The fragment ions derived from electron ionization (70 eV) were detected in a full scan mode (50-450 m/z) and selected ion monitoring mode (74 m/z). The area of the specific ion (m/z 74) was used for quantification of FAMEs.

